# Widespread patterns of sexually dimorphic gene expression in an avian hypothalamic–pituitary–gonadal (HPG) axis

**DOI:** 10.1101/085365

**Authors:** Matthew D. MacManes, Suzanne H. Austin, Andrew Booth, April Austin, Victoria Farrar, Rebecca M. Calisi

## Abstract

The hypothalamic-pituitary-gonadal (HPG) axis is a key biological system required for reproduction and associated sexual behaviors to occur. In the avian reproductive model of the rock dove (*Columba livia*), we characterized the transcript community of each tissue of the HPG axis in both sexes, thereby significantly expanding our mechanistic insight into HPG activity. We report greater sex-biased differential expression in the pituitary as compared to the hypothalamus, with multiple genes more highly expressed in the male pituitary being related to secretory function, and multiple genes more highly expressed in the female pituitary being related to reproduction, growth, and development. We report tissue-specific and sex-biased expression in genes commonly investigated when studying reproduction, highlighting the need for sex parity in future studies. In addition, we uncover new targets of investigation in both sexes, which could potentially change our understanding of HPG function.

## INTRODUCTION

The hypothalamic-pituitary-gonadal (HPG) axis is a system comprised of endocrine glands whose function is vital to the regulation of reproduction and associated behaviors (Fig.1). In all vertebrates studied, from humans to Agnatha, the jawless fishes, the HPG axis is present and its function is generally conserved ^1^. For reproduction to occur, the hypothalamus must produce and secrete gonadotropin-releasing hormone (GnRH), which causes the pituitary gland to secrete gonadotropins, luteinizing hormone (LH) and follicle stimulating hormone (FSH) ^2^. LH and FSH travel through the bloodstream and act upon receptors in the gonads (i.e. testes or ovaries), stimulating gametogenesis and the secretion of sex steroids, such as testosterone and estradiol. These sex steroids then bind with receptors within the HPG axis to create a feedback system that facilitates reproduction and sexual behaviors. The HPG axis has been lauded for providing a foundation to guide reproductive endocrinology investigations, but it has also been criticized as an overly simplistic depiction of the mechanisms mediating reproduction. Now, burgeoning genomic sequencing technologies are permitting a deeper and more complete understanding of the mechanistic drivers of reproduction. They are allowing us to understand how differential gene expression between the sexes could underpin sexually dimorphic reproductive behavior, and how the full complement of genes expressed in the HPG axis could work in concert to define physiology and behavior ^3^.

Previous understanding of HPG function has come about by measuring the amount of circulating hormones in blood, conducting immunochemistry to visualize protein presence or absence, conducting *in situ hybridization* to visualize a target gene, and the use of endocrine and receptor agonists and antagonists and their resulting physiological and behavioral effects. Due to recent technological advances, researchers are beginning to identify correlative and causative links between gene activity and phenotype ^4–10^. Concerning the genomics of reproduction, some studies have generated whole-organism or tissue-specific sequence data of which the HPG axis was a subset ^11,12^. Even fewer studies have reported observing differential gene expression of the HPG axis as a global system ^13^. Here, we focus specifically on patterns of tissue and sexually dimorphic gene expression in the HPG axis of the rock dove (*Columba livia*) and discuss how they might relate to male and female reproductive strategies. Doves have been historically used to study reproductive behavior ^14,15^ and now are proving to be a valuable model for genomics research ^16–18^.

We characterize the first sex-specific avian HPG axis transcriptome using RNA extracted from whole hypothalamus, pituitary, and gonads of sexually mature male and female rock doves (*Columba livia*). We constructed an annotated *de novo* transcriptome assembly for these tissues, which then enabled us to quantify the abundance of all mRNA transcripts per tissue. We elected to perform a *de novo* assembly, rather than use any other alternative approach (*e.g.,* mapping reads to the chicken or rock dove genome), after preliminary mapping experiments suggested that genes important to the behavioral phenotypes in question were not well represented in existing genomic resources. By incorporating candidate gene and *ab initio* approaches across all tissue types and between sexes, we were able to both characterize general patterns of gene expression and identify specific patterns of sex-biased expression. It is our intention that these data illuminate sex differences in gene presence and abundance throughout the HPG axis when birds are at a basal state (i.e. reproductively mature but sampled when not actively breeding). By doing this, we create a resource for the scientific community to devise further studies of how gene expression patterns change over different reproductive stages and in response to physical and social events. As the HPG axis is fairly well-conserved across vertebrates, these data may be useful for formulating potential therapeutic strategies for abnormal HPG axis function.

**Figure 1.**
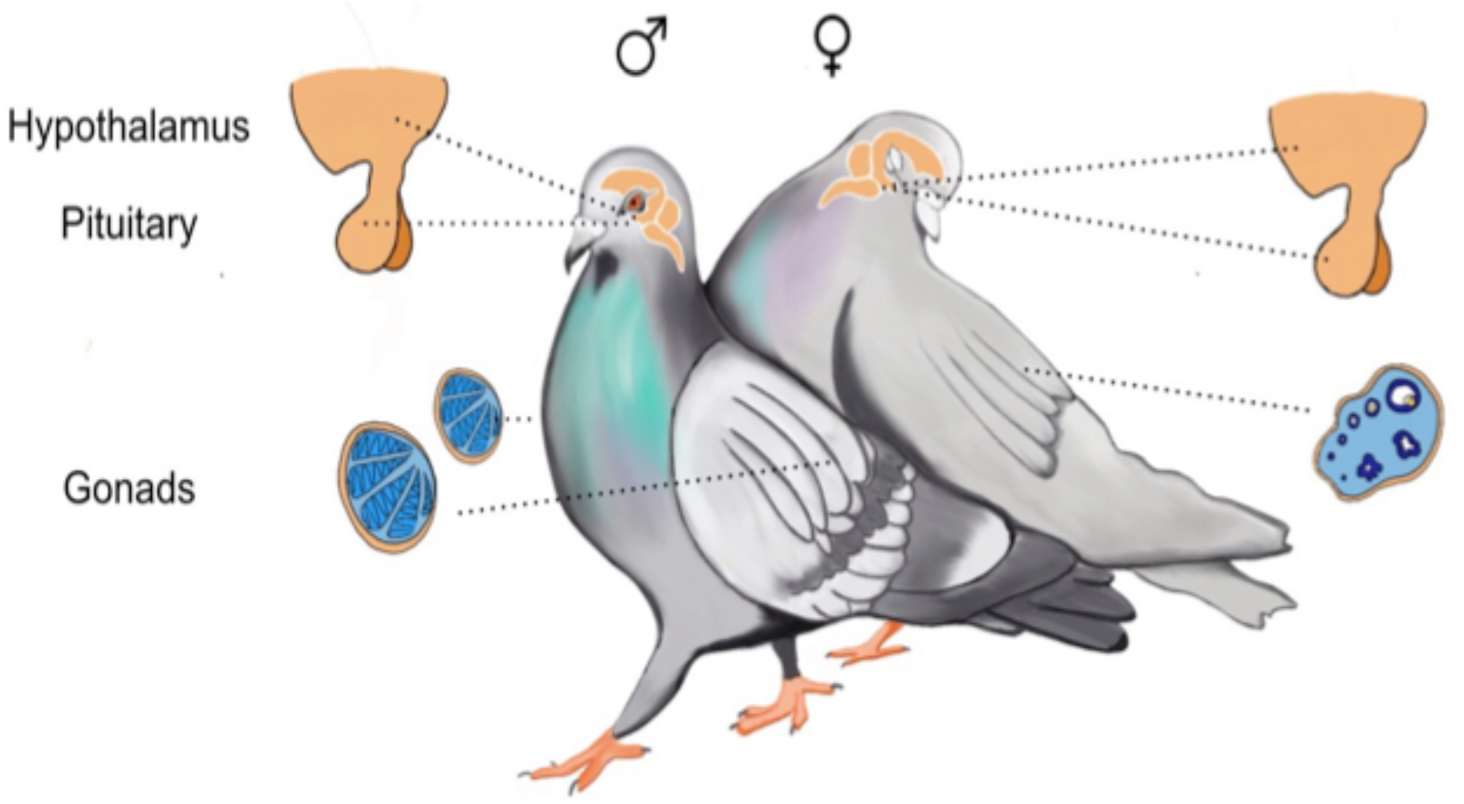
**Depiction of the hypothalamus, pituitary, and gonads of male and female rock doves. Illustration by Natalia Duque.**

## Results

### Sequence Read Data & Code Availability

In total, 20 hypothalamus (11 male, 9 female), 23 pituitary (14 male, 9 female), 13 testes, and 10 ovary samples from 24 birds were sequenced. Each sample was sequenced with between 2.3 million and 24.5 million read pairs. All read data are available using the project ID PRJEB16136. Code used for the analysis of these data are available at https://git.io/vPA09.

### Transcriptome Assembly and Evaluation

The Rock Dove transcriptome assembly consists of 88,011 transcripts (available here https://goo.gl/PYrzas, dryad on acceptance). This assembly, along with the annotations contained in gff3 format, is available at https://goo.gl/DyZ8pw (dryad on acceptance). The transcriptome contains 25,696 with at least 1 hit to the Pfam database, 46,854 hits to OrthoDB^19^, 51,522 hits to the Uniref90 database, 3,108 hits to the transporter database ^20^, and 452 hits to the Rfam database ^21^. These 88,011 assembled transcripts map to 15,102 unique genes in the *Gallus* genome. The evaluation using BUSCO and TransRate are presented in Table 1 and 2 respectively.

**Table 1:** Quality control statistics for the assembled transcriptome: BUSCO metrics include statistics regarding the number of universal vertebrate single copy orthologs found in the assembly. 85.9% of the Avian BUSCOs were identified as full length (complete) sequences, while 4.3% were found to be fragmented. 27.5% were found in greater than 1 copy in the Rock Dove transcriptome.

|  | BUSCO |  |  |
| --- | --- | --- | --- |
|  | Complete | Duplicated | Fragmented |
| Rock Dove 1.0.3 | 85.9% | 27.5% | 4.3% |

**Table 2:** Quality control statistics for the assembled transcriptome: TransRate metrics are derived from mapping RNAseq reads to the assembly, with higher scores indicating a higher quality assembly. A score of 0.41 ranks this assembly higher than the majority of other published transcriptomes, with 90% of reads mapping, and only 2% of bases uncovered (no read support) and 0% contigs low-covered (mean per-base read coverage of < 10).

|  | TRANSRATE |  |  |  |  |  |
| --- | --- | --- | --- | --- | --- | --- |
|  | Score | Number of Contigs | Assembly Size | Percent Mapping | Percent Bases Uncovered | Percent Contigs Lowcovered |
| Rock Dove 1.0.3 | 0.41 | 88011 | 146.8Mb | 90 | 2 | 0 |

### Sequence Read Mapping and Estimation of Gene Expression

Raw sequencing reads corresponding to individual samples of hypothalami, pituitary glands, and gonads were mapped to the Rock Dove reference HPG transcriptome using Salmon, which resulted in between 70% and 80% read mapping. These mapping data were imported into R and summarized into gene-level counts using tximport, after which, edgeR was used to generate normalized estimates of gene expression. Patterns of transcript-expression overlap are presented in Figure 1. Of the 15,102 genes expressed in the HPG, 7,529 are expressed in all tissues (median CPM > 10 - Figure 1). A total of 746 transcripts are expressed in the ovary, hypothalamus, and pituitary, 732 are expressed uniquely in the testes, 529 uniquely in ovary, 418 uniquely in the hypothalamus, and 230 uniquely in the pituitary. Dyads (genes expressed uniquely in pairs of tissues) follow an expected pattern, with the number of genes expressed uniquely in the gonads (testes and ovary) being more than the number of genes expressed uniquely in the pituitary and hypothalamus, followed by the pituitary and ovary, hypothalamus and ovary, pituitary and testes, and hypothalamus and testes. Understanding tissue-specific patterns of expression can provide a glimpse into the HPG axis as an interconnected functional system.

A gene ontology-based pathway analysis^27^ of the transcripts unique to each tissue provides some insights into the specific biological processes important to each tissue. Of the genes that are uniquely expressed in the testes, the three most highly represented pathways include Inflammation mediated by chemokine and cytokine signaling pathway (Panther pathway number P00031), CCKR signaling map (P06959) and Nicotinic acetylcholine receptor signaling pathway (P00044). The top three pathways for genes uniquely expressed in the ovary are related to Gonadotropin-releasing hormone receptor pathway (P06664), Wnt signaling pathway (P00057), and angiogenesis (P00005). Genes uniquely expressed in the hypothalamus cluster around the pathway terms oligodendrocyte differentiation, regulation of glial cell differentiation, positive regulation of synapse assembly, negative regulation of neurogenesis and modulation of synaptic transmission. The pituitary contains uniquely expressed genes related to the Wnt signaling pathway (P00057), the Cadherin signaling pathway (P00012), and the gonadotropin-releasing hormone receptor pathway (P06664).

**Figure 2.**
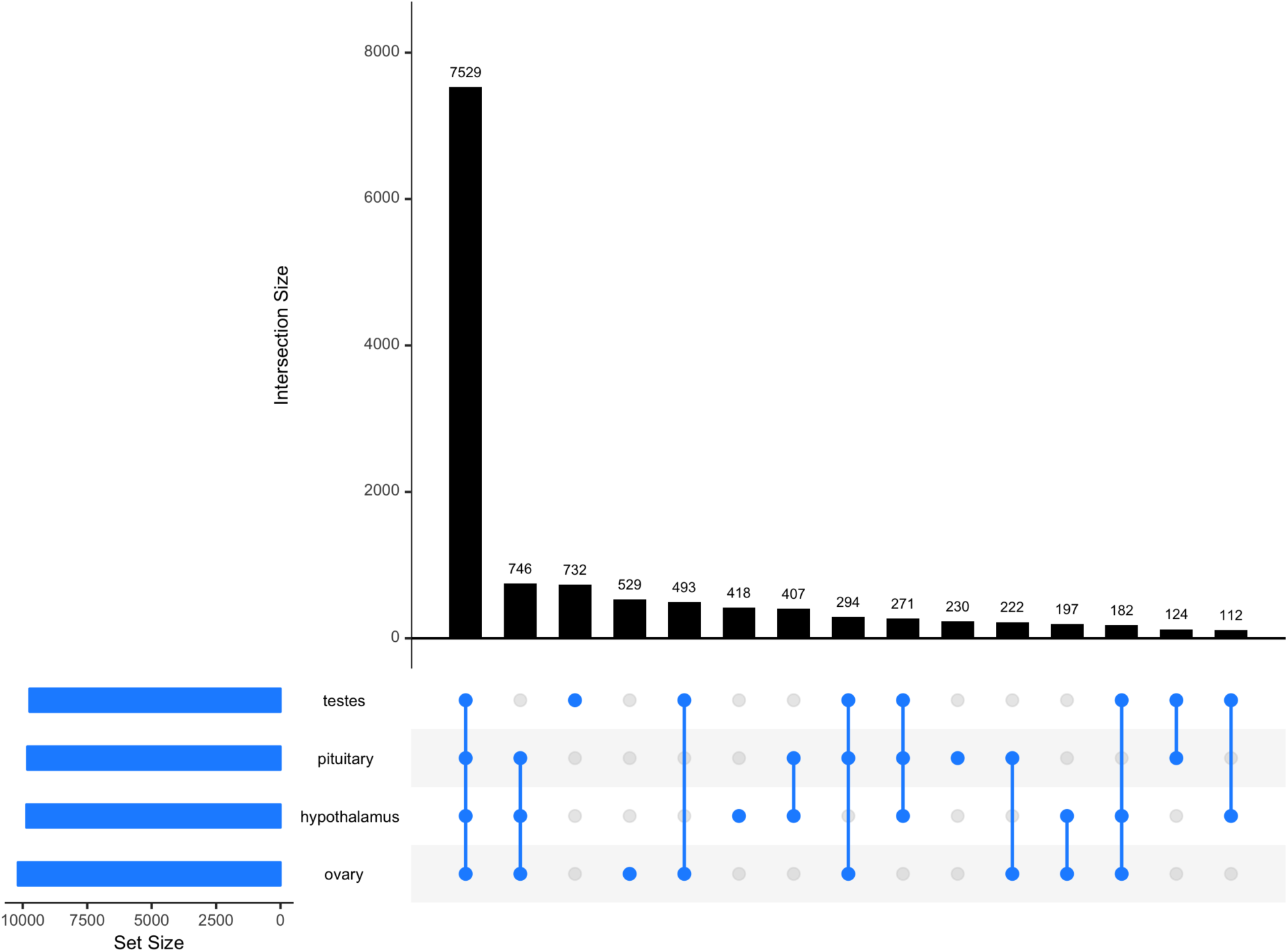
Description of transcript presence overlap in the HPG axis. Bars are annotated by connected dots which indicate a given number of transcripts expressed in those tissues. For example, 746 transcripts are expressed in the pituitary, hypothalamus, and ovary, but these genes are not expressed in the testes. Bars annotated by single blue dots indicate the number of transcripts expressed uniquely in that tissue. For example, 732 transcripts are expressed only in the testes.

### Evaluation of Candidate Gene Expression

Using the assembled transcriptome, we characterized expression in males and females across the HPG axis (e.g., Figure 2). For the candidate genes evaluated, which include gonadotropin releasing hormone (*GnRH−1*) and its receptor (*GnRH−1*−R), LH receptor (*LH−R*), FSH receptor (*FSH−R*), androgen receptor (*AR*), estrogen receptor alpha and beta (*ESR alpha*, *ESR beta*). In addition, we examined gonadotropin inhibitory hormone (*GnIH*), due to its inhibitory effect on the HPG axis and reproductive behavior ^22^, vasoactive intestinal peptide (*VIP*), due to its role in stimulating the release of the pituitary hormone, prolactin ^23^, and its receptor (*VIP−R*), prolactin (PRL) due to its facilitation of parental care behaviors, including crop milk production ^24^, and its receptor (*PRL−R*), arginine vasopressin-like receptor 1A and 1B (*AVPR1A, AVPR1B*), mesotocin (*MT*) and its receptor (*MTR*), due to their role in social bonding behaviors ^25,26^, progesterone receptor (*PGR*) due to its role in reproductive behavior ^27^, CYP19, which encodes the aromatase enzyme (*ARO*), which is responsible for the conversion of testosterone into estradiol ^28^.

Using a generalized linear model and least-squares means for post-hoc tests of significance (P<0.05), we uncovered greater expression of *GnRH-1-R*, *AR*, and *PGR* in the female pituitary as compared to the male pituitary (z-ratio 2.324, p-value =0.0201; z ratio = −3.842, p-value = 0.0001; z ratio = 5.559, p-value = <.0001, respectively). We also found greater expression of *PRL* in the female pituitary (z ratio = 2.912, p-value = 0.0036) as compared to the male pituitary, but *PRL* was more highly expressed in the male hypothalamus as compared to the female hypothalamus (z ratio = −2.433, p-value = 0.0150). The *PRL* receptor was also more highly expressed in the male hypothalamus and pituitary gland as compared to corresponding female tissues (z ratio −2.674, p-value = 0.0075 and z ratio −2.564, p-value = 0.0104, respectively). Finally, we discovered greater expression of *AVPR1A* in the male pituitary as compared to the female pituitary (z ratio = −2.416, p-value = 0.0157).

**Figure 2A-F.**
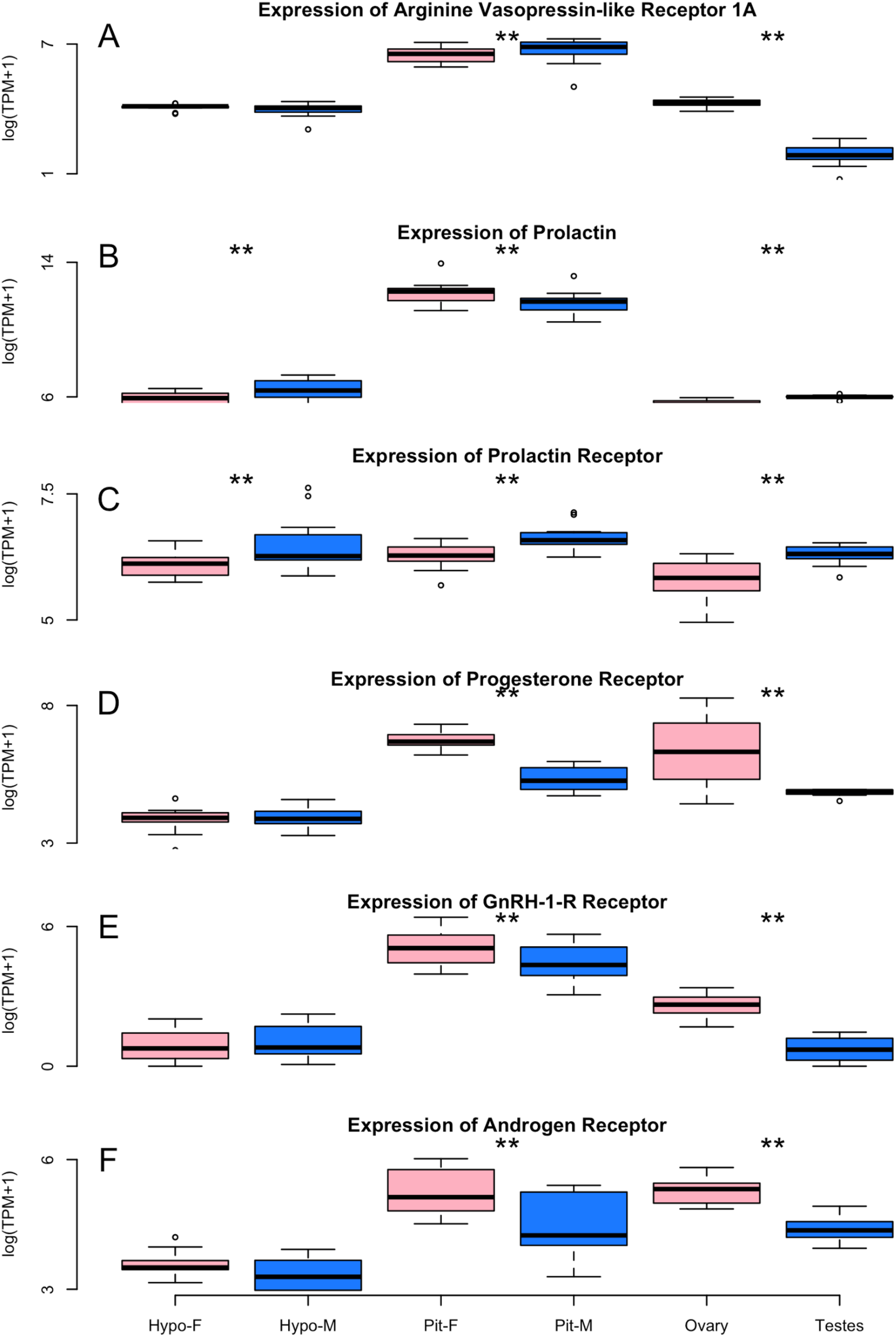
Box and whisker plots illustrating the differential expression between the sexes of six candidate genes. The Y-axis indicates levels of gene expression as presented by taking the log of transcripts per million plus one (TPM+1) while tissue and sex are presented on the X-axis. The bottom and top of the box indicates the first and third quartiles. The band inside the box is the median. Whiskers indicate the 95% confidence intervals of the data. ** denotes statistically significant differences in expression.

**Table 3:** Mean gene expression (normalized counts per million (CPM)) +/− standard deviation) for a list of candidate genes known to be important mediators of reproductive biology.

|  | Hypo |  | Pit |  | Gonads |  |
| --- | --- | --- | --- | --- | --- | --- |
| Gene | Female | Male | Female | Male | Female | Male |
| *GnRH-I* | 116.1 +/- 75.9 | 142.3 +/- 121.6 | 0.3 +/- 0.5 | 1.3 +/- 2.3 | 0.6 +/- 0.7 | 1.2 +/- 1.1 |
| *GnRH-I-R* | 2.34 +/- 2.67 | 2.85 +/- 2.75 | 245.6 +/- 212.4 | 112.8 +/- 83.76 | 14.0 +/- 6.7 | 1.38 +/- 1.09 |
| *LH-R* | 19.0 +/- 11.3 | 23.03 +/- 12.2 | 0.9 +/- 0.95 | 0.38 +/- 0.67 | 264.45 +/- 77.54 | 42.59 +/- 39.1 |
| *FSH-R* | 4.02 +/- 4.42 | 5.13 +/- 2.62 | 9.4 +/- 3.75 | 11.5 +/- 5.62 | 151.59 +/- 80.2 | 181.0 +/- 51.56 |
| *AR* | 37.7 +/- 13.4 | 27.8 +/- 13.1 | 226.6 +/- 121.4 | 111.1 +/- 72.7 | 204.1 +/- 62.5 | 80.3 +/- 23.1 |
| *ERa* | 49.2 +/- 14.0 | 57.2 +/- 18.6 | 724.5 +/- 329.3 | 836 +/- 283.4 | 166.2 +/- 152.5 | 64.6 +/- 32.6 |
| *ERb* | 41.9 +/- 18.7 | 36.1 +/- 14.5 | 53.8 +/- 16.5 | 58.6 +/- 27.9 | 173.4 +/- 90.5 | 7.4 +/- 3.9 |
| *GnIH* | 287.8 +/- 148.7 | 383.2 +/- 510.0 | 4.0 +/- 3.8 | 2.9 +/- 3.1 | 0.1 +/- 0.2 | 0.7 +/- 0.7 |
| *VIP* | 1.3e+02 +/- 60.1 | 1.3e+02 +/- 98.3 | 1.19 +/- 1.37 | .46 +/- .77 | 1.3e+01 +/- 15.6 | 0 +/- 0 |
| *VIP-R* | 70.78 +/- 18.5 | 69.2 +/- 24.5 | 47.3 +/- 141.5 | 376.35 +/- 134.64 | 42.3 +/- 17.3 | 18.8 +/- 7.9 |
| *PRL* | 389.9 +/- 170.4 | 766.0 +/- 436.7 | 295840.2 +/- 320026.2 | 136607.5 +/- 121634.2 | 250.9 +/- 95.2 | 397.9 +/- 45.1 |
| *PRL-R* | 463.0 +/- 141.7 | 803.4 +/- 558.4 | 540.4 +/- 143.2 | 790.85 +/- 204 | 338.5 +/- 131.8 | 550.66 +/- 100.7 |
| *AVPR1A* | 117.3 +/- 58.8 | 98.3 +/- 34.4 | 202.2 +/- 111.3 | 362.2 +/- 181.6 | 19.0 +/- 13.5 | 3.2 +/- 2.3 |
| *AVPR1B* | 5.4 +/- 5.4 | 5.4 +/- 3.1 | 1538.3 +/- 503.5 | 1900.1 +/- 709.1 | 3.8 +/- 1.3 | 2.3 +/- 1.5 |
| *MT* | 264.9 +/- 82.1 | 206.9 +/- 101.1 | 173 +/- 39.9 | 172.78 +/- 60.9 | 182.11 +/- 265.4 | 11.1 +/- 2.98 |
| *MTR* | <1 | <1 | <1 | <1 | <1 | <1 |
| *PGR* | 50.6 +/- 24.6 | 51.4 +/- 20.8 | 874.4 +/- 294.9 | 216.7 +/- 82.6 | 1004.6 +/- 1193 | 126.7 +/- 14.4 |
| *CYP19* | 212.6 +/- 137.5 | 325.8 +/- 200.2 | 26.4 +/- 10.3 | 49.5 +/- 20.2 | 786.6 +/- 446.2 | 1.7 +/- 1.3 |

### Global Evaluation of Gene Expression

Analysis of global patterns of gene expression, aimed at uncovering previously unknown sex-specific differences in gene expression in the hypothalamus and pituitary by examining the entire transcriptome, was conducted using edgeR. After controlling for over 15,000 comparisons, we normalized the count data using the TMM method ^29^, which is done by finding a set of scaling factors for the library sizes that minimize the log-fold changes between the samples for most genes. This analysis revealed sex-specific differences in gene expression, particularly in the pituitary, where 218 genes were more highly expressed in males, while 153 genes were more highly expressed in females (Figure 3). The five most differentially expressed genes are Zinc Finger AN1-Type Containing 5 (*ZFAND5*), Betacellulin (*BTC*) which appears to be responsive to LH ^30^, Cbp/P300 Interacting Transactivator With Glu/Asp Rich Carboxy-Terminal Domain 4 (*CITED4*), Growth Regulation By Estrogen In Breast Cancer 1 (*GREB1*) which is regulated by estrogen ^31^, and Kinesin Family Member 24 (*KIF24*). All had false discovery rates (FDR) < 7e-09. The full table of differentially expressed genes is available at https://git.io/vXJWp, which includes several tantalizing novel candidates, including Ecto-NOX Disulfide-Thiol Exchanger 1 (*ENOX1*; Fig. 4). In contrast to the pituitary, the hypothalamus exhibited only a single differentially expressed gene. That gene was potassium voltage-gated channel subfamily Q member 1 (*KCNQ1*, FDR =0.007). KCNQ1, which we found to be more highly expressed in females, is known to be imprinted (expressed in a parent-of-origin-specific manner) in mammals ^32^.

A number of expression patterns in the pituitary were identified by gene ontology analysis, which uses a controlled vocabulary to describe gene function and relationships between these terms. First, gene ontology terms that describe genes differentially expressed in the female pituitary (Table 4) are clustered around terms related to the binding of lipid-based hormones, immune function, growth, and development. Gene ontology terms from genes differentially expressed in male pituitary (Table 5) are clustered around terms related to muscles and movement.

**Figure 3.**
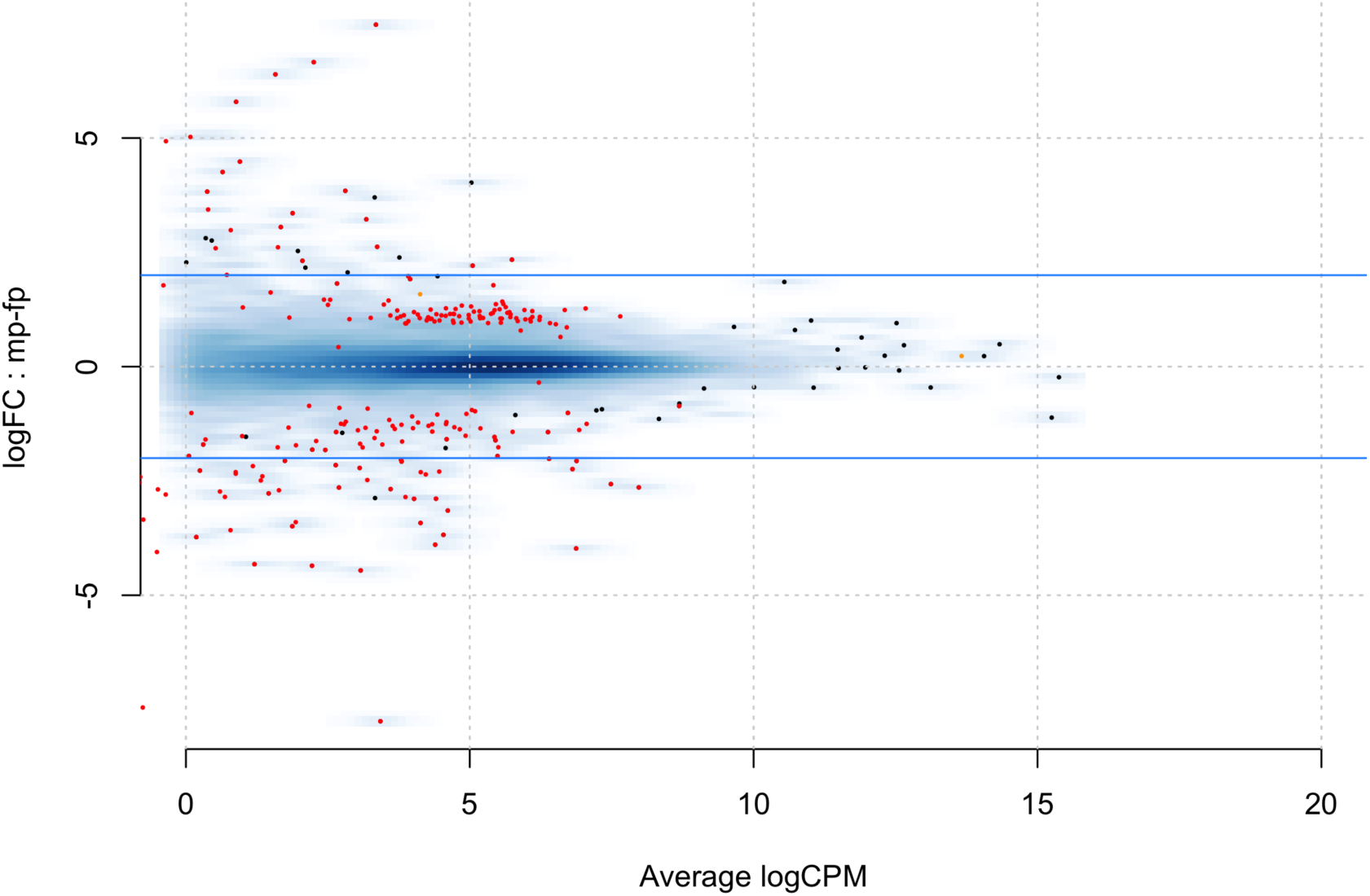
MA plot for the *ab initio* comparison between genes differentially expressed in the male and female pituitary. Red dots indicate statistically different genes. Those with positive values on the log fold change (logFC) Y axis are more highly expressed in the male pituitary, while those with negative values are more highly expressed in the female pituitary. The x-axis is a measure of gene expression, with higher numbers indicating higher levels of gene expression.

**Figure 4.**
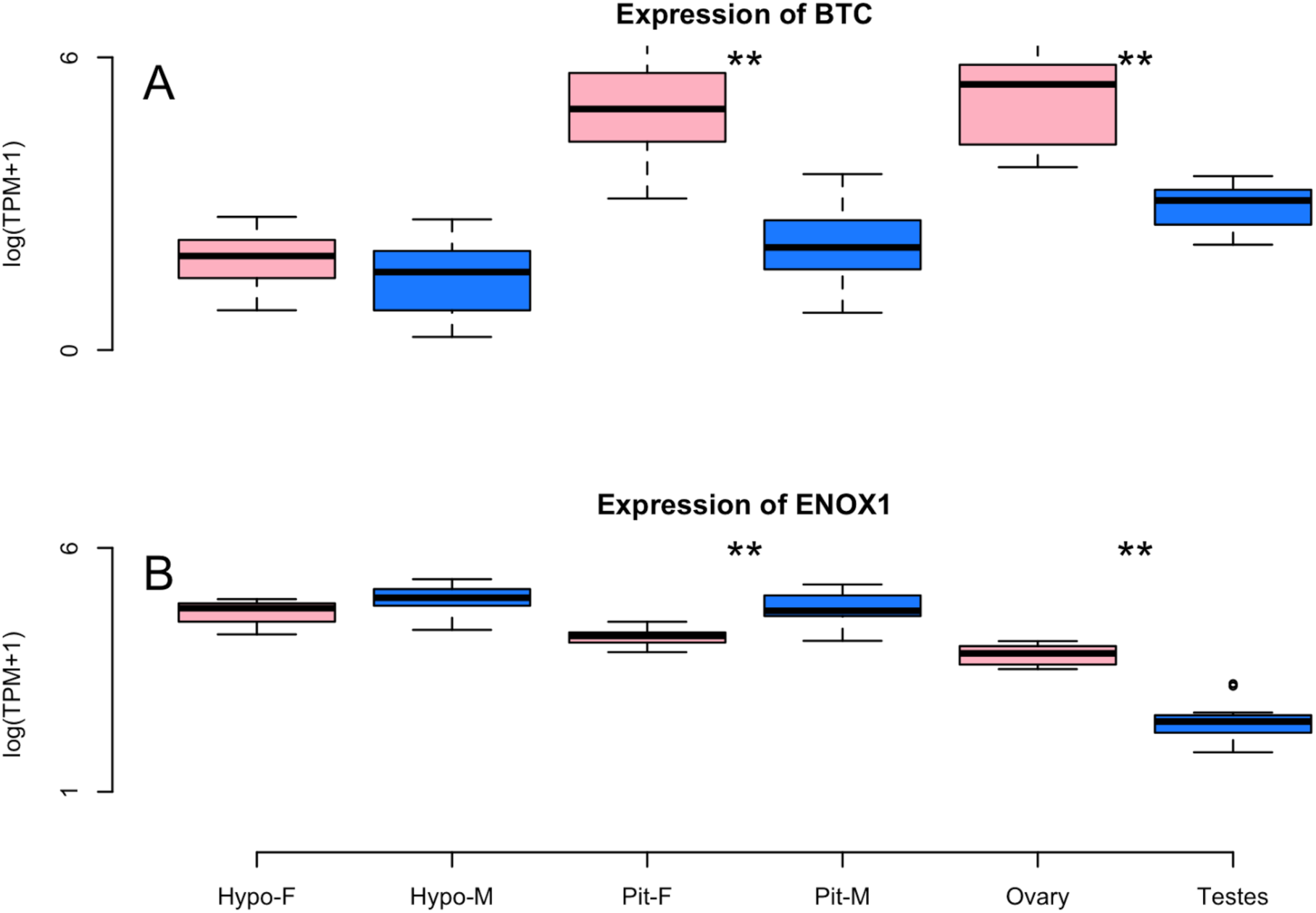
Box and whisker plots illustrating sex-biased expression of two genes, Betacellulin (*BTC*) and Ecto-NOX Disulfide-Thiol Exchanger 1 (*ENOX1*), identified using our global analysis. The Y-axis indicates levels of gene expression as presented by taking the log of transcripts per million plus one (TPM+1) while tissue and sex are presented on the X-axis. The bottom and top of the box indicates the first and third quartiles. The band inside the box is the median. Whiskers indicate the 95% confidence intervals of the data. ** denotes statistically significant differences in expression.

**Table 4:** **Top 20 Gene Ontology (GO) Terms** describing differentially expressed genes in the female pituitary gland. These terms describe the putative function of genes using a controlled vocabulary. Abbreviations: Ontology (Ont), Molecular Function (MF), Biological Processes (BP). N signifies the number of genes in the entire dataset that are linked to the specific ontology term. F+ indicates the number of differentially expressed genes significantly more highly expressed in the female pituitary, while M+ indicates the number of expressed genes significantly more highly expressed in the male pituitary. Results are limited to terms where the specific term contains >4 genes. In all cases, p-values are corrected for multiple hypothesis tests using the Bonferroni correction.

| GO number | Term | Ont | N | F+ | M+ | P-value |
| --- | --- | --- | --- | --- | --- | --- |
| **GO:0008289** | lipid binding | MF | 50 | 6 | 1 | 1.92E-05 |
| **GO:0048469** | cell maturation | BP | 13 | 3 | 0 | 4.29E-04 |
| **GO:0021700** | developmental maturation | BP | 18 | 3 | 0 | 1.18E-03 |
| **GO:0043178** | alcohol binding | MF | 7 | 2 | 0 | 2.93E-03 |
| **GO:0033293** | monocarboxylic acid binding | MF | 8 | 2 | 0 | 3.88E-03 |
| **GO:0014066** | regulation of phosphatidylinositol 3-kinase signaling | BP | 8 | 2 | 0 | 3.88E-03 |
| **GO:0006869** | lipid transport | BP | 27 | 3 | 2 | 3.92E-03 |
| **GO:0007548** | sex differentiation | BP | 27 | 3 | 0 | 3.92E-03 |
| **GO:0010876** | lipid localization | BP | 29 | 3 | 2 | 4.82E-03 |
| **GO:0048017** | inositol lipid-mediated signaling | BP | 9 | 2 | 0 | 4.95E-03 |
| **GO:0014065** | phosphatidylinositol 3-kinase signaling | BP | 9 | 2 | 0 | 4.95E-03 |
| **GO:0048015** | phosphatidylinositol-mediated signaling | BP | 9 | 2 | 0 | 4.95E-03 |
| **GO:0005496** | steroid binding | MF | 9 | 2 | 0 | 4.95E-03 |
| **GO:0005319** | lipid transporter activity | MF | 10 | 2 | 1 | 6.14E-03 |
| **GO:0043627** | response to estrogen | BP | 10 | 2 | 0 | 6.14E-03 |
| **GO:0004867** | serine-type endopeptidase inhibitor activity | MF | 12 | 2 | 0 | 8.86E-03 |
| **GO:0042698** | ovulation cycle | BP | 13 | 2 | 0 | 1.04E-02 |
| **GO:0022602** | ovulation cycle process | BP | 13 | 2 | 0 | 1.04E-02 |
| **GO:0005102** | receptor binding | MF | 114 | 5 | 0 | 1.07E-02 |
| **GO:0046545** | development of primary female sexual characteristics | BP | 14 | 2 | 0 | 1.20E-02 |

**Table 5:** **Gene Ontology (GO) Terms** describing genes differentially expressed genes in the male pituitary gland. These terms describe the putative function of genes using a controlled vocabulary. Abbreviations: Ont= Ontology (Ont), MF (Molecular Function (MF), BP (Biological Processes (BP). N signifies= the number of genes in the entire dataset that are linked to the specific ontology term. F+ indicates the number of differentially expressed genes significantly more highly expressed in the female pituitary, while M+ indicates the number of expressed genes significantly more highly expressed in the male pituitary. Results are limited to terms where the specific term contains >4 genes. In all cases, p-values are corrected for multiple hypothesis tests using the Bonferroni correction.

| GO number | Term | Ont | N | F+ | M+ | P-value |
| --- | --- | --- | --- | --- | --- | --- |
| **GO:0006936** | muscle contraction | BP | 21 | 0 | 4 | 3.90E-04 |
| **GO:0006941** | striated muscle contraction | BP | 10 | 0 | 3 | 5.53E-04 |
| **GO:0003012** | muscle system process | BP | 26 | 0 | 4 | 9.15E-04 |
| **GO:0050879** | multicellular organismal movement | BP | 4 | 0 | 2 | 1.77E-03 |
| **GO:0050881** | musculoskeletal movement | BP | 4 | 0 | 2 | 1.77E-03 |
| **GO:0003009** | skeletal muscle contraction | BP | 4 | 0 | 2 | 1.77E-03 |
| **GO:0008092** | cytoskeletal protein binding | MF | 101 | 1 | 6 | 7.20E-03 |
| **GO:0003779** | actin binding | MF | 53 | 0 | 4 | 1.28E-02 |

## Discussion

The HPG axis is a system comprised of endocrine tissues whose function is vital to the regulation of reproduction and associated behavior. Here, we report patterns of gene expression amongst tissues of the HPG axis as well as patterns of sex-biased gene expression within them. We describe patterns of tissue-specific expression and shared incidence of expression between tissues, providing an important picture of the functional-connectivity of these tissues. We also describe sex-biased patterns of gene expression for a set of candidate genes, currently known to play important roles in reproduction and associated behaviors. Lastly, we use an *ab initio* approach to identify all differentially expressed transcripts between male and female pituitary and hypothalamus. Our findings provide a vital resource for researchers examining the molecular basis of reproductive behavior and the mechanisms underlying fundamental physiological and behavioral differences in males and females.

### The Columba livia HPG transcriptome

We present an HPG transcriptome that is substantially complete, with fewer than 10% of universal avian orthologs missing (Table 1). This number suggests that all, or nearly all, transcripts present in the avian HPG have been successfully reconstructed, and the missing avian orthologs may be expressed in tissues other than those studied here. The transcriptome is structurally sound, as is evident from its TransRate score of.41, with 90% of reads mapping concordantly to the assembled reference (Table 2).

### Evaluation of Candidate Gene Expression

Decades of previous research describing the HPG axis and its role in reproductive biology have elucidated much of the molecular machinery underlying phenotypes ^33^. Based on this previous work, we targeted well-known substrates involved in the facilitation and mediation of reproductive processes and their associated behaviors for investigation (Table 3). Other genes not highlighted here can be found at https://git.io/vXJW.

We discovered previously unrecognized sex-specific differences in several of these candidate genes. Arginine vasopressin-like receptor 1A (*AVPR1A*) is known to be implicated in reproductive behaviors such as pair-bonding and parental care ^34–37^. *AVPR1A* is most highly expressed in the pituitary, followed by the ovary, the hypothalamus, and lastly the testes. We found statistically significant differences in expression of this gene in the pituitary, with expression of this gene to be 1.8x higher in males as compared to females. Although a small number of studies have investigated sex-specific differences in gene expression ^38–40^, including one in which expression in males was linked to pair-bond formation ^41^, it is not well known if *AVPR1A* is differentially expressed by sex. As fields of behavior and genomics further integrate, links between sex-specific roles in pair-bonding and reproductive behavior in the context of corresponding differences in gene expression will transform the way we understand how behavior is regulated.

We also investigated other candidate genes involved in reproduction and associated behaviors. Prolactin (*PRL*) is an important promoter of parental care, promoting lactation ^42^ and related to appetite and weight gain ^43^. We found *PRL* to be more highly expressed in the male hypothalamus as compared to the female hypothalamus. The receptor for prolactin (*PRL-R*) was also more highly expressed in the male hypothalamus and pituitary as compared to females. In contrast, *PRL* was more highly expressed in the female pituitary as compared to the male pituitary. Though this species exhibits bi-parental care, including the production of crop milk by both males and females, neither sex was caring for eggs or chicks at the time of collection to help control for potential reproductive stage confounds. However, the interesting sexually-biased differences we note in inter-tissue gene expression inspires the question as to whether *PRL* is regulated in a sex-specific manner to produce a sex-specific result? Or, despite the difference in location of expression, does *PRL* activation lead to sexually monomorphic prolactin-mediated reproductive processes and behaviors? In other words, the differences observed in underlying genetic expression between the sexes may converge to the same behavioral endpoint, permitting certain factors in one sex to offset the effects in the other, making the sexes more similar ^44^. Post our discovery that *PRL* expression differs relative to sex and tissue of the HPG axis, the next step would be to investigate how the action of *PRL* on each of these tissues manifests in males versus females.

In addition, we found *GnRH-1-R*, *AR*, and *PRG* to be more highly expressed in the female pituitary as compared to the male pituitary. GnRH-1, produced in the hypothalamus, is a major mediator of gonadotropin release in the pituitary. These gonadotropins signal the gonads to produce androgens, which feedback onto androgen receptors along the HPG axis. We found that androgen receptors are more highly expressed in females as compared to males. Females tend to have lower circulation of androgens than males, so could this be a way to increase sensitivity to androgen signals in females? Finally, progesterone receptor is associated with the mediation of female breeding cycles ^45^, but it can also serve as an important precursor to the creation of multiple hormones in males and females, including testosterone and estradiol. We found that the androgen receptor is more highly expressed in the female pituitary relative to the male pituitary. While expression in both sexes has been linked to reproductive development ^46–48^, to our knowledge, sex differences in expression have never been reported. To better elucidate the phenomenon of these sexually-dimorphic genetic expression phenotypes, further studies of functional genomics are required. Indeed, study of the most well-characterized candidate genes underlying reproductive behavior could reveal important and previously unrecognized sex-differences in expression. As the field of behavioral genomics advances, important questions to ask will be, are these differences biologically relevant, and if so, how do differences in expression lead to differences in behavior and reproductive output? Future work focused on these relationships will significantly shape our understanding of the molecular mechanisms underlying the observed patterns.

### Global Evaluation of Gene Expression

Avian reproductive behavior has been shown to be heavily influenced by the endocrine function of the hypothalamus, pituitary, and gonads. Although sex-biased differences in vertebrate reproductive behavior have been well-noted, data on sex-specific patterns of gene expression that could be influencing these behaviors are lacking. This may largely be due to the novelty of various genomic technologies and the expense of collecting such data. However, it is becoming increasingly feasible and affordable to conduct such studies, making possible the potential for a more integrative understanding of reproductive behavior ^3^. Thus, in addition to investigating sexually dimorphic candidate gene expression in the HPG axis, we used an *ab initio* RNAseq approach to explore differences in *all* sexually dimorphic HPG gene expression in each tissue. The newly discovered sex-biased genetic differences that we report have the potential to offer a more in-depth picture of sexual dimorphism at the molecular level.

To conduct a global analysis of gene expression, we compared levels of expression in the tissues of the HPG axis shared by both sexes, the pituitary glands and hypothalami. Unlike our previous analyses where we targeted genes of interest, here we analyzed all genes expressed, controlling for over 15,000 multiple comparisons. We uncovered 361 sex-based differentially-expressed genes in the pituitary: 218 were more highly expressed in males, while 153 were more highly expressed in females. A table detailing differences in expression of all expressed genes, including those statistically and non-statistically differentially expressed, can be accessed at https://git.io/vXJWp. In contrast, only a single gene, the potassium voltage-gated channel subfamily Q member 1 (*KCNQ1*), was statistically differentially expressed in the hypothalamus (full table available here https://git.io/vXJW5. This profound disparity in patterns of differential expression between hypothalamic and pituitary tissues begs a question of function - how do the different tissues of the HPG axis contribute to physiological and behavioral differences between males and females? Is the pituitary more involved in the maintenance of sex-specific physiology and behavior than the hypothalamus? Studies comparing gene expression in males and females are rare - most often studies have combined the sexes (but see ^38–40,49–54^). However, Nishida and colleagues ^39^ found similar patterns of gene expression in mice, reporting that 43 genes were more highly expressed in the female pituitary while only 3 were more highly expressed in the hypothalamus as compared to males. Could this pattern of sex-biased expression be conserved across vertebrates, and if so, can we conclude that the pituitary, more so than the hypothalamus, plays a greater role in the maintenance of sex-specific reproductive physiology and behavior?

Another hypothesis as to why we may observe less differential expression at the level of the hypothalamus is because of the high heterogeneity of the region, which could result in the dilution of observable differences in gene expression existing in more discrete locations. On the other hand, high heterogeneity of the hypothalamus might only mask results of differential expression if individual nuclei express antiparallel sex-specific changes, resulting in an averaging of gene expression signals. Given this particular scenario is likely to be exceptionally rare, it is possible that nuclei heterogeneity of the hypothalamus does not contribute significantly to the observed patterns, though future investigations may yield more telling data if specific nuclei are examined within the hypothalamus.

Gene ontology (GO) analysis of the differential expression in the pituitary revealed interesting patterns of enrichment, or patterns of gene expression related to a common biological function or process. GO terms enriched for genes expressed more highly in males were often related to motor or muscle function, with specific terms such as muscle contraction being more highly expressed in males as compared to females. The differentially expressed genes linked to these GO terms include *Myosin* and other components of the myosin actin skeleton. Previous research has identified a potential role for myosin in the pituitary, including in its primary function of secretion ^55–57^. For example, GnRH-induced secretion of LH can be altered by manipulating the action of myosin in the anterior pituitary ^56^. In females, a surge in LH will trigger ovulation. We were unable to control for the specific ovulatory stage of the females sampled, though by chance we can assume that most were not sampled at the exact point they were experiencing their LH surge. However, males were sexually mature and actively paired with female partners at the time of sampling. In males, LH stimulates Leydig cell production of testosterone. Thus, one explanation for having GO term enrichment for genes more highly expressed in males under this label of motor control is that the male pituitary could be more sensitive to the GnRH signal to stimulate LH release due to myosin-related actions. The biological relevance of sex-specific differential expression of *Myosin* remains unknown and deserves future study.

Gene ontology terms enriched for genes expressed more highly in the female pituitary were often related to aspects of female growth and development. This may be due to their varying stages of follicular development. Females, unlike males, must grow and maintain follicles and the oviduct, which is energetically costly ^59^. In a passerine species, females increased their resting metabolic rate by 22% during egg laying, of which 18% was attributed to maintenance of the oviduct ^59^. While birds in our study were not actively nesting, we could not control for reproductive stage of the gonads. When we extracted female gonads, we found that they were experiencing different stages of follicular development. Some birds had regressed follicles, but many had a follicular hierarchy. The presence of follicular hierarchy could explain the presence of gene ontologies related to ovulation. Similarly, lipid binding and transport may be associated with follicular development because females are laying down lipid-rich yolk. On the other hand, males never had regressed testes. The active state of the male gonads is typical of pigeons who breed year-round, except for a brief photorefractory period in the North temperate fall where they may regress ^60^. Gonads of females are more plastic and vary with the presence of suitable mates and breeding stage ^60^.

Using the global analysis approach, we were able to identify sex-biased differentially expressed genes that have not previously been targeted for investigation of HPG function. Such findings that are yielded from this discovery-based approach offer promising targets for future study, inspiring new lines of investigation ^3^. One particularly promising target for understanding sex differences in reproductive biology is Betacellulin (*BTC*), which is highly expressed in the female pituitary as compared to the male pituitary (4.4 FC - Figure 4A). *BTC* is an epidermal-like growth hormone known to regulate female reproduction ^61^ via mediation of LH ^30^ and regulation of progesterone receptors ^62^. In addition to being differentially expressed in the pituitary, it is also highly expressed in the female ovary, but not the hypothalamus. It is lowly expressed, but present in all male tissues, which begs the question, what role could *BTC* have in male reproductive physiology, if any?

The Ecto-NOX Disulfide-Thiol Exchanger 1 (*ENOX1*) gene, an electron transport gene, is more highly expressed in the male pituitary relative to the female pituitary (1.2x FC - Figure 4B). This gene, known to be involved in circadian rhythms ^63^, has also been associated with weight gain ^64^, litter size in the pig ^65^, and eggshell thickness in a genome-wide association study (GWAS) of chickens ^66^. What role this gene plays in the pituitary, and specifically in the male pituitary, where it is differentially expressed, is completely unclear. Further work aimed at understanding this and the other differentially expressed genes can be the key to gaining the most complete understanding yet of how the HPG axis functions.

## Summary

Researchers have described many important characteristics of the molecular basis for reproductive behaviors, including the characterization of multiple genes expressed in the HPG axis that are critical to reproduction. Our sex-specific, global analyses have uncovered hundreds of sexually-biased genes. These genes can provide novel clues to assist behavioral biologists and functional genomicists in more fully understanding the mechanisms underlying sexual dimorphism in reproductive behaviors. Indeed, future work can now use this information to better understand causal relationships and functions of genes expressed differently in males and females.

In conclusion, we reveal patterns of tissue specific and sexually dimorphic gene expression in the HPG axis. We report sex-biased expression in genes commonly investigated when studying reproduction. In addition, we offer up promising new targets of investigation that could lead to a better understanding of HPG function in both sexes. Our results highlight the need for sex parity in transcriptomic studies, providing new lines of investigation of the mechanisms of reproductive function.

## Methods

### Animal Collection Methods

Birds were housed at the University of California, Davis, in large, outdoor aviaries (5’×4’×7’), with 8 sexually reproductive adult pairs per aviary. All birds were three years old and had been housed in their respective aviaries for one year at the time of collection. All birds were also successful breeders, having paired naturally and produced multiple clutches throughout the year thus far. However, to control for reproductive stage as much as was possible, all birds used in this study were sampled at a time when they were naturally without eggs or chicks, creating a type of “baseline” sampling point. All sampling occurred during late fall of 2015 and winter of 2016 between 10:00-12:00 (PST) to avoid potential seasonal and circadian rhythm confounds. While birds were exposed to natural light, we also augmented their aviaries with artificial lights set to 14L:10D photoperiod in order to maintain reproductive condition. All methods were carried out in accordance with relevant guidelines and regulations, and all experimental protocols were approved by the University of California, Davis, IACUC (#18895). Fourteen males and ten females were sacrificed within 3 min of entering their aviary. They were anesthetized using isoflurane prior to decapitation. Brains, pituitaries, and gonads were extracted and immediately placed on dry ice and transferred within the hour to a −80°C freezer until further processing. Frozen brains were coronally sectioned on a cryostat (Leica CM 1860) at 100µM and the hypothalamus was isolated using surgical punches. We used Karten and Hodos’ stereotaxic atlas ^67^ of the brain of the pigeon to locate the hypothalamus and collect it in its entirety. In brief, we, collected hypothalamic tissue beginning at the point of bifurcation of the tractus septomesencephalicus and ending after the cerebellum was well apparent. Hypothalamic, pituitary, and gonadal tissue were preserved at −80°C in RNALater and shipped from the University of California, Davis, to the University of New Hampshire for further processing.

### Library Preparation and Sequencing

Tissues frozen in RNALater were thawed on ice in an RNAse-free work environment. Total RNA was extracted using a standard Trizol extraction protocol (Thermo Fisher Scientific, Waltham, MA). The quality of the resultant extracted total RNA was characterized using the Tapestation 2200 instrument (Agilent, Santa Clara, CA), after which Illumina sequence libraries were prepared using the TruSeq RNA Stranded LT Kit (Illumina). The Tapestation was, again, used to determine the quality and concentration of these libraries. Each library was diluted to 2nM with sterile ddH_2_O, and pooled in a multiplexed library sample. The multiplexed library sample was then sent to the New York Genome Center for 125 or 150 base pair paired-end sequencing on a HiSeq 2500 platform.

### Sequence Quality Control and Assembly

Sequence read data were downloaded and quality checked using FastQC version 0.11.5 ^68^. A *de novo* transcriptome for the HPG was assembled following the Oyster River Protocol for Transcriptome Assembly ^69^. In brief, 59 million paired end reads from two individuals (1 male and 1 female) were error corrected using RCorrector version 1.0.2 ^70^. Adapters, as well as bases with a Phred score <2 were trimmed using Skewer 0.2.2 ^71^, and assembly was carried out using Trinity version 2.2.0 ^72^, Binpacker version 1.0 ^73^, and Shannon version 0.0.2 ^74^, after which the resultant transcriptomes were merged using the software package transfuse version 0.5.0 (https://github.com/cboursnell/transfuse). Lowly expressed transcripts, defined as those with an abundance of less than 0.5 transcript-per-million (TPM<0.5), were filtered out of the dataset. The resultant assembly was annotated using the software package dammit (https://github.com/camillescott/dammit), and evaluated using BUSCO 2.0 beta 3 ^75^ and TransRate version 1.0.1 ^76^.

### Mapping and Differential Gene Expression

All samples were used (20 hypothalamus (11male, 9 female), 23 pituitary (14 male, 9 female), 13 testes, and 10 ovary samples from 24 birds.) Raw reads were mapped to the annotated transcriptome after a quasi-mapping index was prepared using Salmon 0.7.0 ^77^. Rock dove transcripts were mapped to genes from the *Gallus gallus* genome version 5, using BLAST ^78^. All data were then imported into the R statistical package (version 3.3.0) ^79^ using tximport ^80^ for gene level evaluation of gene expression, which was calculated using edgeR (version 3.1.4) ^81^ following TMM normalization and correction for multiple hypothesis tests by setting the false discovery rate (FDR) to 1%.

### Gene Expression Evaluation

Although excellent genomic resources exist for the Rock Dove ^17^<u>, the current Rock Dove genome available lacks functional annotation (*e.g.,* gene ontology terms, information about function and protein networks).</u> Thus, we elected to establish orthologous relationships between transcript sequences we generated in Rock Dove with those from the *Gallus gallus* genome, version 5.

Patterns of gene expression were evaluated using two distinct methods. First, we selected *apriori* genes of interest based on their known involvement in reproduction and associated behaviors (Table 3). Differences in gene expression were evaluated between these genes in the hypothalamic, pituitary, and gonadal tissues and between both sexes using a generalized linear model framework (expression ~ sex * tissue) with post-hoc significance for all pairwise combinations of factors tested using the Bioconductor package lsmeans (https://cran.r-project.org/package=lsmeans). Second, a more global investigation of differential expression was conducted to characterize the general presence and expression of all genes per tissue and how they might differ between the sexes. A comparison of differential expression between tissues was not carried out secondary to results from preliminary analysis suggesting that most genes are expressed differently, thus, making meaningful comparisons uninformative. Instead, to characterize patterns of expression in each tissue, we generated a transcript presence/absence matrix (1=present, 0=absent) using median CPM (counts-per-million, generated in edgeR) > 10 as the metric. Overlaps were visualized using the R package UpSet ^82^. Gene ontology enrichment analysis was carried out using the Kolmogorov-Smirnov test ^83^ for significance in the R package, topGO ^84^.

## Acknowledgements

This work is supported by NSF IOS 1455960 to RMC and MM. We thank Jesse Krause, Jonathan Peréz, and members of the Calisi Lab for their help in maintaining the rock dove aviaries and aiding in tissue collection. We also thank Stacia Sower, Samuel Díaz-Muñoz, and the University of California, Davis, Environmental Endocrinology Group (EEG), comprised of the labs of Rebecca Calisi, Tom Hahn, Marilyn Ramenofsky, Karen Ryan, and John Wingfield, for their comments on this project and manuscript. We are grateful to Natalia Duque for her illustration (Fig. 1).

## Author Contribution Statements

Matthew MacManes and Andrew Lang constructed sequencing libraries, did bioinformatics analyses, and wrote the paper. Suzanne H. Austin, April Booth, Victoria Farrar and Rebecca Calisi conducted live animal experiments and wrote the paper.

## Additional Information

Read data is available at PRJEB16136

